# Engineering of Transmembrane Alkane Monooxygenases to Improve a Key Reaction Step in the Synthesis of Polymer Precursor Tulipalin A

**DOI:** 10.1101/2024.07.04.601532

**Authors:** Andrea Nigl, Veronica Delsoglio, Marina Grgić, Lenny Malihan-Yap, Kamela Myrtollari, Jelena Spasic, Margit Winkler, Gustav Oberdorfer, Andreas Taden, Iva Anić, Robert Kourist

## Abstract

The α-methylene-γ-butyrolactone tulipalin A, naturally found in tulips can polymerize via addition at the vinyl group or via ring-opening polymerization, making it a highly promising monomer for biobased polymers. As tulipalin A biosynthesis in plants remains elusive, we propose a pathway for its synthesis starting from the metabolic intermediate isoprenol. For this, terminal hydroxylation of the α-methylene substrate isoprenyl acetate is a decisive step. While a panel of fungal unspecific peroxygenases showed a preference for the undesired epoxidation of the *exo-*olefin group, bacterial alkane monooxygenases were specific for terminal hydroxylation. A combination of protein engineering based on *de novo* structure prediction of the membrane enzymes with cell engineering allowed to increase the specific activity by 6-fold to 1.83 U g_cdw_ ^-1^, unlocking this reaction for the fermentative production of tulipalin A from renewable resources.

---

Tulipalin A (**6**, α-methylene-γ-butyrolactone) is a natural secondary metabolite found in plants among the Alstroemeriaceae and Liliaceae families. It shows strong antimicrobial and insecticidal properties.^[1–3]^ **6** combines two functional moieties accessible for polymerization within one molecule – a lactone and a vinyl group. Its *exo*-methylene group makes tulipalin A a potential substitute for the petrol-based monomer methyl methacrylate (MMA), an important component for reactive adhesives, coatings, and sealants.^[4]^ The lactone can also undergo sustainable ring-opening polymerization (ROP) resulting in unsaturated (bio-degradable) polyesters that can be further cross-linked resulting in polymeric networks.^[4–7]^ The versatility of tulipalin A makes it an interesting building block for a broad range of high-performance polymers.^[4,6]^ While tulipalin A can be extracted from tulips, isolating a bulk chemical from plant material is not economically feasible. Unfortunately, the biosynthesis *in planta* remains mostly unknown, representing a major obstacle to the development of biotechnological processes.^[8–11]^ Herein, we propose synthesis of **6** starting from isoprenol **1** (**Scheme 1A**), which can be readily produced in engineered microorganisms from the terpenoid precursor isopentenyl pyrophosphate (IPP) either via the mevalonate (MVA) or the methyl-erythritol-phosphate (MEP) pathway.^[12–15]^ We suggest linking the synthesis of isoprenol **1** to the production of tulipalin A **6** via an initial acylation step to protect the C1-OH moiety of isoprenol from oxidation in the later phase of the pathway, followed by selective hydroxylation of isoprenyl acetate **2** at the C4-position. The formed alcohol **3a** can then easily be oxidized either enzymatically^[16,17]^ or chemically^[18]^ to the carboxylic acid **5**, which can be directly lactonized to tulipalin A **6** by acidification at elevated temperatures (**Figure S9**).^[10,18]^

The regioselective hydroxylation of the *O-*protected isoprenol (isoprenyl acetate **2**) to 4-acetoxy-2-methylene-butan-1-ol **3a** is the crucial step in this synthetic route. It requires a chemo- and regioselective oxygenase that neither catalyzes the epoxidation of the *exo*-double bond to **3b** nor the internal hydroxylation at the C2-atom to **3c** (**Scheme 1B**).

**Scheme 1.**
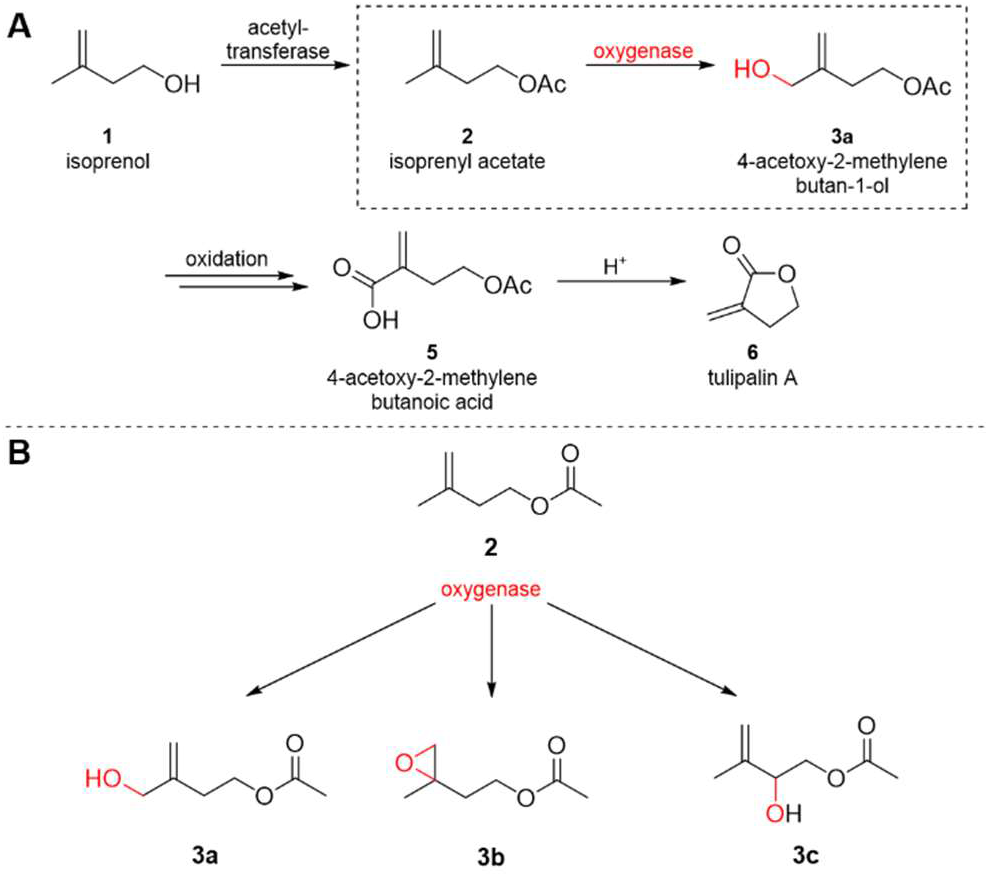
**(A)** Artificial downstream pathway towards tulipalin A deviating from the hemiterpenoid isoprenol **1. (B)** Identification of an oxygenase selective for terminal hydroxylation of **2** at the C4-position to **3a** without epoxidation to **3b** and/ or internal hydroxylation to **3c** is key in this synthetic route.

Since the discovery of unspecific peroxygenases (UPO) from *Agrocybe aegerita* (*Aae*UPO), these highly active and stable enzymes have attracted increasing attention for cell-free C-H-oxyfunctionalization reactions, making them promising candidates for the desired hydroxylation of **2**. Investigation of a commercially available enzyme panel of 77 UPOs revealed that *Aae*UPO and 71 other UPOs indeed accepted **2**. Unfortunately, the formation of **3a** was not observed for any of the enzymes within this panel (**Figure S3, Table S7**). Instead, the epoxide **3b** (**Figure 1A**) was formed.^[19,20]^ Due to the strong preferences of the screened UPOs for epoxidation, we expected it to be unlikely to identify a peroxygenase selective for C4-hydroxylation. A complete switch of chemoselectivity from epoxidation to hydroxylation by enzyme engineering appeared to be exceedingly difficult.

**Figure 1.**
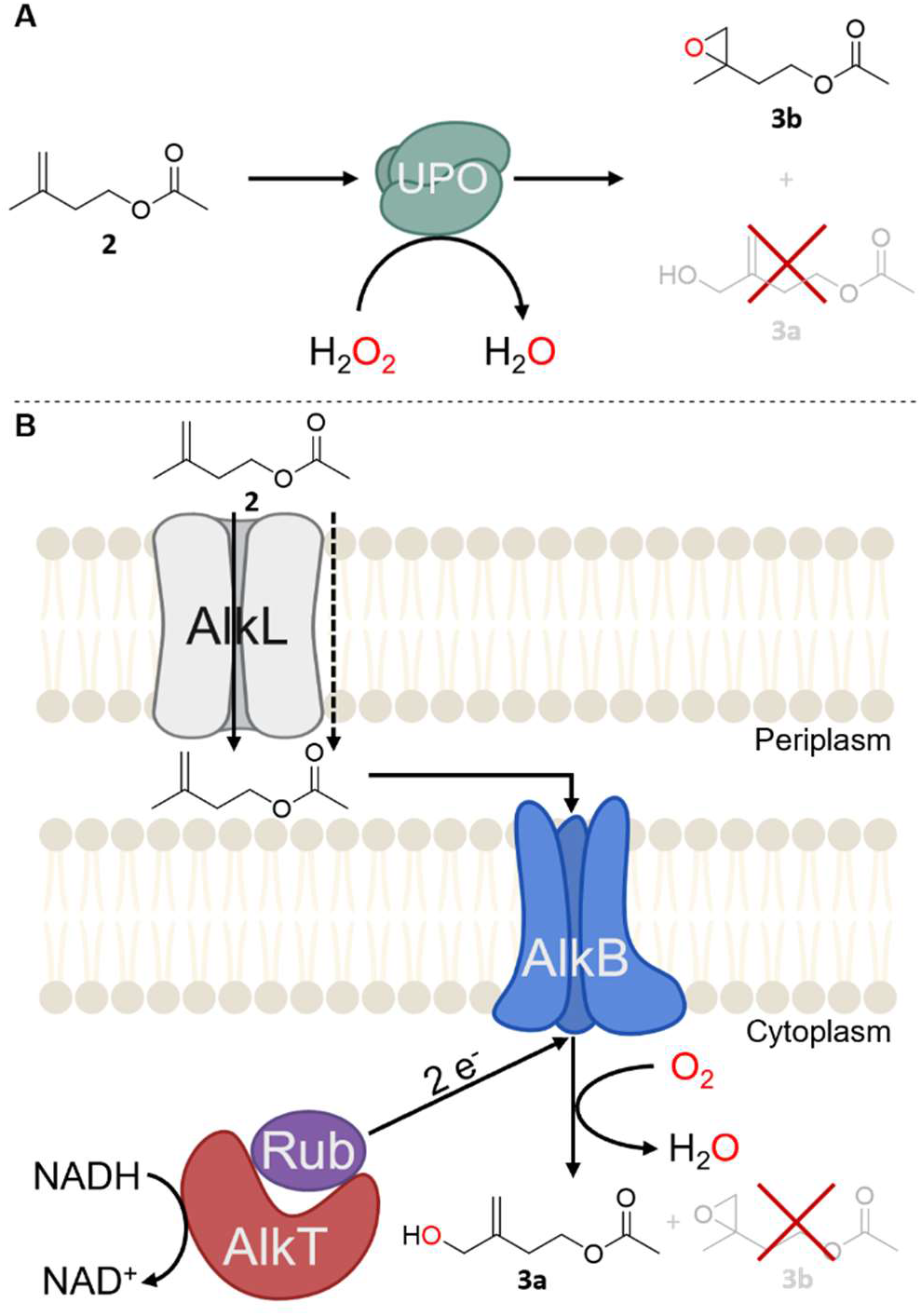
Conversion of isoprenyl acetate **2** by heme-dependent UPOs gives rise to the epoxide **3b (A)** while the reaction with the AlkBFGT(L) yields the desired terminal alcohol **3a (B)**.

Hence, we investigated a different type of monooxygenases for the selective hydroxylation of **2**: The well-known alkane monooxygenase (MO) AlkB from *Pseudomonas putida* GPo1 (PpGPo1AlkB) has been reported to catalyze the hydroxylation of the similar isobutene to 2-methyl-2-propen-1-ol while leaving the *exo*-methylene group unreacted.^[21]^ The natural function of the transmembrane diiron non-heme hydroxylase AlkB system encoded by the *alk*-operon is the initial hydroxylation reaction of an alkane degrading pathway present in many soil and marine bacteria.^[22,23]^ AlkB shows remarkable selectivity for ω-hydroxylation of linear medium-chain alkanes. This is a striking difference to the widely applied CYP450 monooxygenases that often hydroxylate at the ω-1 and ω-2 positions of linear alkanes and medium-chain fatty acids esters.^[24–29]^ Furthermore, AlkB was shown to hydroxylate the acetate esters of various fatty alcohols without acting on the acetate group.^[30]^ As an aside, the successful application of the AlkB system for the ω-hydroxylation of fatty acid methyl esters at an industrial scale underlines the industrial applicability of the system.^[31–33]^ As PpGPo1AlkB was reported to convert the sterically hindered isobutene to 2-methyl-2-propen-1-ol^[21]^ we envisioned selective hydroxylation of **2** without undesired oxidation of the double bond or internal hydroxylation.

In this study, the genes of the AlkBFGT system from *P. putida* GPo1 were recombinantly expressed in *Escherichia coli* BL21 (DE3) under the control of the natural alkane-inducible *alkB* promoter.^[27,34]^ GC-MS analysis of whole-cell biotransformations with **2** indicated a product peak with a mass spectrogram fitting to the desired product 4-acetoxy-2-methylene-butan-1-ol **3a**. For verification with NMR-spectroscopy, the reaction was scaled up to 20 mL in shake flasks, the product was extracted from the cell suspension and purified via column chromatography. The recorded data agreed with literature-reported spectra for **3a** (^1^H NMR ((300 MHz, CDCl_3_), *δ* 5.12 (s, 1H), 4.94 (s, 1H), 4.22 (t, *J* = 6.72 Hz, 2H), 4.11 (s, 2H), 2.42 (t, *J* = 6.39 Hz, 2H), 2.05 (s, 3H) ppm).^[35]^ Formation of the epoxide **3b** or the internal hydroxylation was not observed (**Figure 1B**).

Besides chemo- and regioselectivity, the specific activity of the biocatalyst is a decisive parameter for industrial application. The activity of *E. coli* BL21 (DE3) PpGPo1AlkBFGT in the whole-cell biotransformation was 0.28 U g_CDW_ ^-1^. While this allowed the identification of **3a**, the activity of AlkB with the sterically hindered **2** was significantly lower compared to the natural alkane substrate *n-*octane (4.43 U g_CDW_ ^-1^) and deemed to be a bottleneck for the application in the artificial biosynthesis route (**Figure 2**). To increase the product formation rate in the whole-cell biotransformations, three different strategies were employed: a) exploration of homologous enzymes for their ability to hydroxylate **2;** b) use of the outer-membrane transporter AlkL to facilitate cellular uptake, and c) semi-rational engineering of PpGPo1AlkB based on literature data^[36,37]^, *de novo* structure prediction ^[38]^ and substrate docking.^[27,32]^

**Figure 2.**
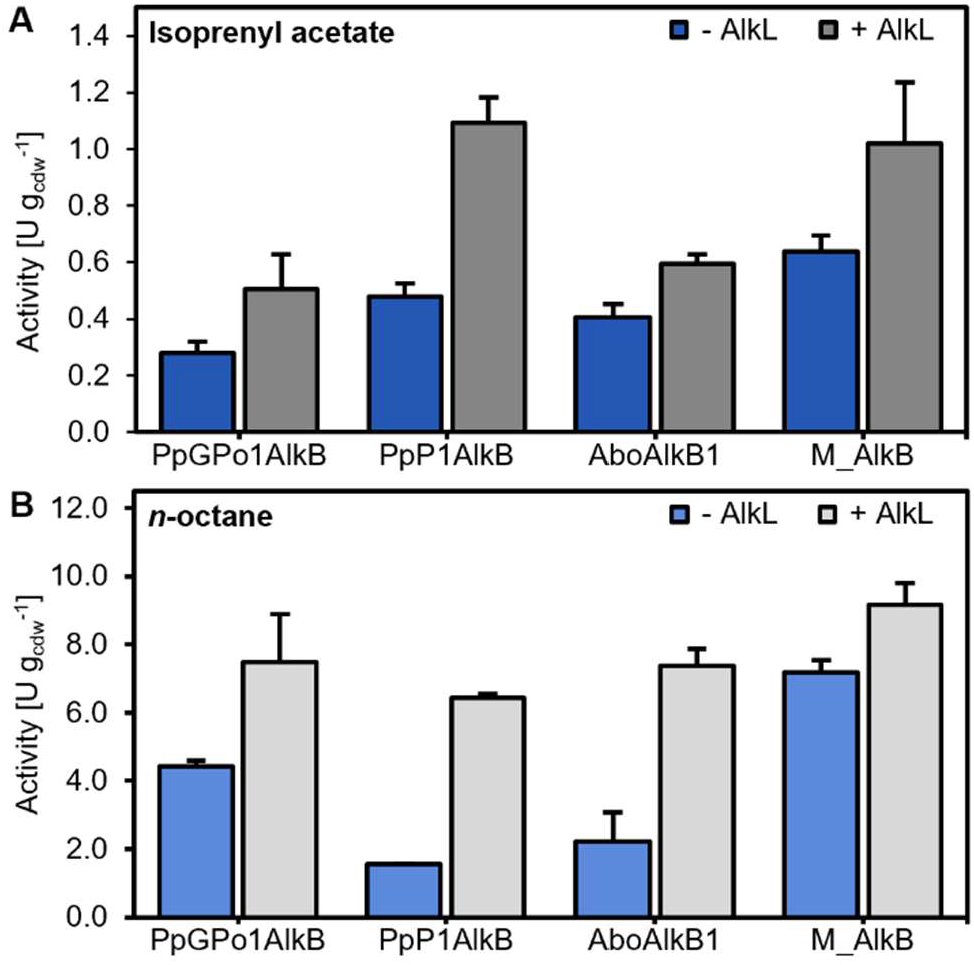
Activities of AlkB homologs from different organisms towards the target substrate isoprenyl acetate **2 (A)** and *n*-octane **(B)**. The presence of the outer-membrane transporter AlkL led to improved substrate uptake, thereby increasing the overall activity of the whole-cell system.

A set of four homologous alkane MOs with varying sequence identities to PpGPo1AlkB were selected. AlkB from *P. putida* P1 (PpP1AlkB; 90 % sequence identity) and *Alcanivorax borkumensis* (AboAlkB1; 77 %) were described in the literature to have a similar substrate scope as PpGPo1AlkB and were shown to accept electrons from the PpGPo1 rubredoxins.^[39,40]^ The alkane monooxygenase from *Acinetobacter baylyi* (AbaAlkB; 41 %) is known to act on medium to long-chain alkanes and was selected as a more distantly related enzyme.^[41]^ The enzyme from *Marinobacter* sp. originates from a marine metagenome study and was annotated as putative alkane MO (M_AlkB; 78 %).^[42]^ The genes were cloned into the same expression vector replacing the gene encoding the *PpGPo1AlkB* while retaining the genes encoding the electron transport system of *P. putida* GPo1. The functionality of the enzymes was tested using *n-*octane as a reference substrate (**Figure 2B**). As shown in **Figure 2A** PpP1AlkB and AboAlkB1 were similarly active towards isoprenyl acetate, while AbaAlkB was poorly produced (data not shown), and no activity was observed. This enzyme was therefore not further investigated. PpP1AlkB, AboAlkB1, and M_AlkB showed a similar substrate preference as PpGPo1AlkB when comparing the activity with **2** and *n-*octane. M_AlkB from *Marinobacter* sp. was found to be about 2-fold more active towards both substrates. No internal hydroxylation nor epoxide formation with **2** as substrate was observed with either of the alkane monooxygenases. This underlined the outstanding selectivity of these enzymes and confirmed our initial expectation based on the reactivity of AlkB towards isobutene.^[21]^

Facilitated transport of hydrophobic substrates across the outer cell membrane is an important factor in whole-cell biocatalysis. The co-expression of the gene *alkL* encoding for an outer-membrane transporter protein was shown to increase the uptake of hydrophobic substrates thereby increasing the overall activity.^[27,32]^ While AlkL is crucial for the efficient transport, particularly of larger substrates (>C10), we expected it might as well facilitate the cellular uptake of the shorter but sterically hindered **2**. Therefore, we expanded the *alkB(homolog)FGT* operon by the *alkL* gene. In **Figure 2A** and **2B**, a clear trend of increased whole-cell activity in the presence of AlkL can be seen for **2** (up to 2-fold) and *n*-octane (up to 4-fold), respectively.

Enzyme engineering of membrane-bound enzymes remains a challenge: Limited knowledge of their structure and reaction mechanisms represents an obstacle to their rational engineering.^[25,26,36]^ In lieu of accurate structural information on AlkB, random mutagenesis has been the method of choice thus far. In two epPCR-based directed evolution studies of PpGPo1AlkB, the selection of improved variants via alkane utilization revealed that substituting the bulky W55 to a smaller serine shifted the substrate preference to alkanes ≥ C_12_. The triple mutant V129M+L132V+I233V showed higher activities towards short-chain alkanes (C_4_-C_5_).^[36,37]^ These variants presented themselves as starting points for the optimization of the biocatalyst. Recently, two cryo-EM structures of AlkB from *Fontimonas thermophila* FtAlkB were published (pdb code: 8SBB and 8F6T, respectively.) sharing about 60 % sequence identity with PpGPo1AlkB, allowing the selection of amino acid residues based on structural considerations.^[28,29]^ The PpGPo1AlkB single variants W55S, V129M, L132V, I233V, as well as the triple variant V129M+L132V+I233V (MVV) and quadruple variant MVV+W55S (SMVV) were tested in whole-cell biotransformations with **2** and *n*-octane as substrates (**Figure 3**).^[36,37]^ In addition, a docking study was conducted based on the predicted AlphaFold2 (AF2) structure of PpGPo1AlkB.^[38]^ The two Fe-atoms in the catalytic center, essential for the enzymatic activity and coordinated by four highly conserved motifs containing nine histidines, were modeled into the AF2-structure (**Figure 4**).^[26,28,29,36,43]^ Isoprenyl acetate **2** was then docked into the inner catalytic pocket of PpGPo1AlkB. Residues lining the binding pocket exhibiting high energies and closest to the best docking pose were subjected to computational mutagenesis analysis using Rosetta FastDesign, whereby the sampled amino acid identities were restricted to hydrophobic amino acids only.^[44]^ The selected BP residues (N135, F164, L265, A268 and L309) in the active site are shown in **Figure 4**. At position L309 no other energetically more favorable amino acid than Leu was identified during the Rosetta redesign. The two most frequently found single mutants for N135, F164, L265, and A268 were experimentally characterized. Investigation of these variants (**Figure 3A** and **B**) in whole-cell biotransformations with **2** and *n-* octane, respectively, revealed that the mutation I233V seems to be an activity driver for isoprenyl acetate and *n-*octane. Interestingly, the mutation F164L resulted in higher activities towards **2** but decreased the activity towards the natural substrate. Substituting F164 to amino acids similar to leucine (I and V) lowered the activity for both substrates.

**Figure 3.**
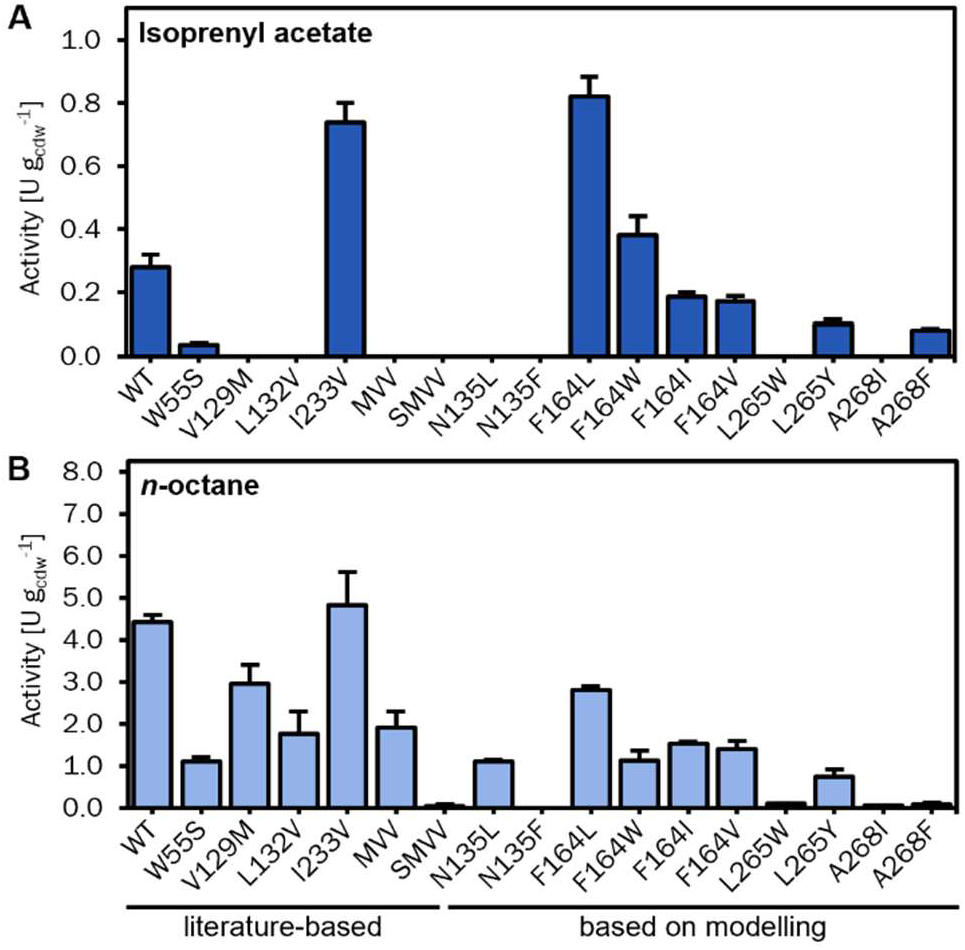
Activity screening of PpGPo1AlkB variants in whole-cell bio-transformations towards the target substrate **2 (A)** and the natural substrate *n-*octane **(B)**. The library was generated by site-directed mutagenesis based on previously reported variants and computational modeling based on the AF2-structure. The activity towards *n-*octane is represented as overall activity including octanal and octanoic acid as the products of overoxidation.

**Figure 4.**
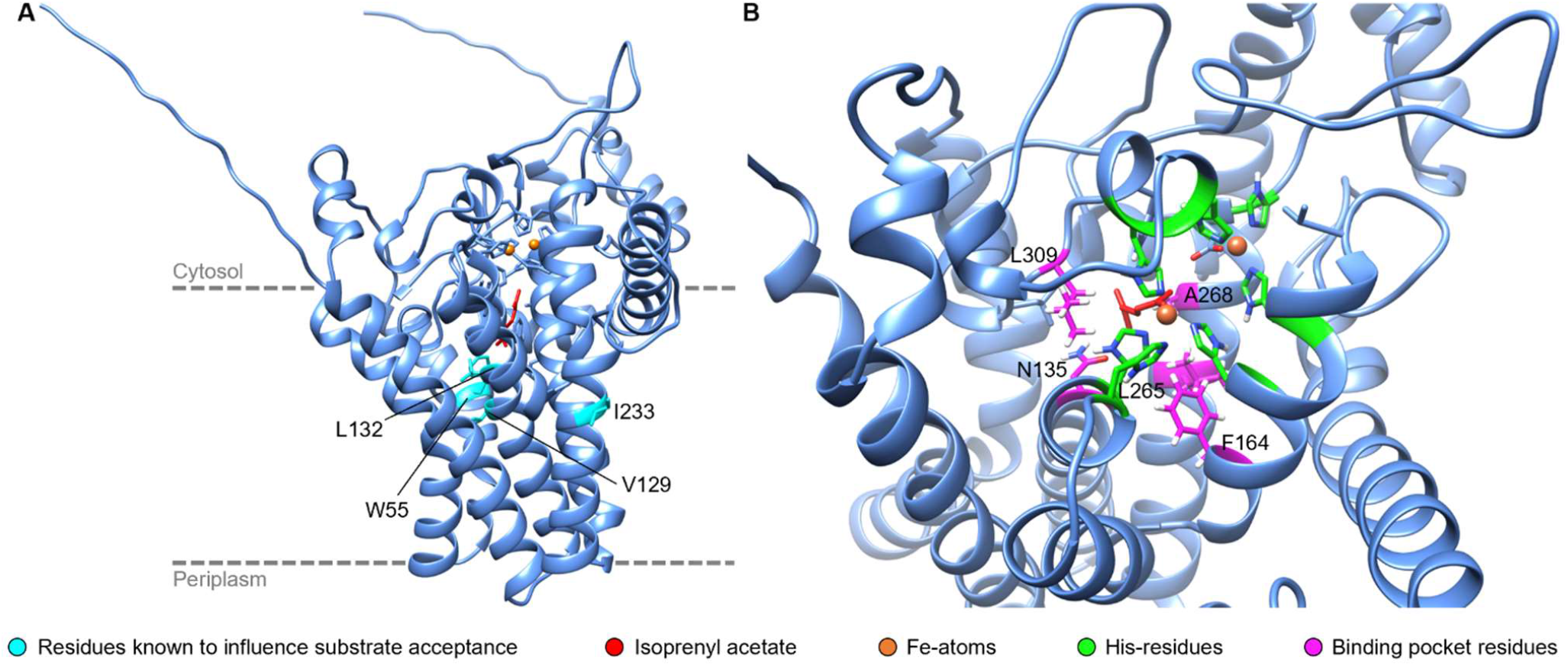
**(A)** Design of PpGPo1AlkB based on the AF2-model with the iron atoms (orange spheres) coordinated by highly conserved histidines in the catalytic center used for docking studies with isoprenyl acetate **2** (red) and the prediction of active site mutations. The residues reported to influence substrate acceptance in AlkB (W55, V129, L132, and I233) are highlighted in cyan. The dashed grey line approximates the position of the enzyme in the inner cell membrane. **(B)** Binding pocket of PpGPo1AlkB. Residues close to the docked substrate and showing high Rosetta Energy Units were selected as targets for mutation studies: N135, F164, L265, A268, and L309 are shown in pink.

We sought to investigate the potential enhancement of the formation of **3a** by combining the different engineering approaches in PpGPo1AlkB. Surprisingly, no additive effect of the double mutation I233V+F164L was observed (**Table 1**; activity represents **3a** formation only). As expected, the co-expression of *alkL* resulted in higher activities with I233V, F164L and the combined variant. As M_AlkB already exhibits increased activity towards **2**, we wanted to see if the substitutions I238V and F169L to this monooxygenase in combination with AlkL would lead to a similar boost, as shown for PpGPo1AlkB (**Table 1**). Indeed, the I238V and F169L variants were also more active towards **2** than the WT. Similar to PpGPo1AlkB, no additive effect of the combined mutations (F169L + I238V) in M_AlkB was observed.

**Table 1.** Specific activities and relative activities (variant over WT) of improved PpGPo1AlkB and M_AlkB WT and variants (with AlkL only) towards **2**. The substrate preference *P* represents the ratio of the activity with **2** over *n-*octane, while the preference shift compares the change in preference of the variant over the WT enzyme.

| Enzyme | Variant (V) | Isoprenyl acetate | | Substrate preference *P*<br>$v_2/v_{n-octane}$ | Preference shift<br>$P_V/P_{WT}$ |
| --- | --- | --- | --- | --- | --- |
|  |  | Specific activity [U g<sub>cdw</sub><sup>-1</sup>] | Relative activity |  |  |
| PpGPo1AlkB | WT | 0.51 ± 0.12 | 1 | 0.1 | - |
|  | I233V<sup>[37]</sup> | 0.81 ± 0.28 | 1.61 | 0.1 | 2.1 |
|  | F164L | 1.08 ± 0.13 | 2.1 | 0.2 | 3.1 |
|  | I233V+F164L | 0.87 ± 0.19 | 1.71 | 0.1 | 2.0 |
| M_AlkB | WT | 1.02 ± 0.22 | 1 | 0.1 | - |
|  | I238V | 1.83 ± 0.32 | 1.79 | 0.2 | 2.1 |
|  | F169L | 0.91 ± 0.03 | 0.89 | 0.2 | 1.6 |
|  | I238V+F169L | 0.63 ± 0.01 | 0.62 | 0.1 | 1.1 |

In this work, we demonstrate for the first time the improvement of the activity of AlkB MOs towards an unnatural sterically hindered substrate by enzyme engineering. The striking selectivity of AlkB for terminal hydroxylation of **2** to **3a** is crucial for its application in the synthesis leading to tulipalin A **6**. On the contrary, UPO-catalyzed reactions with **2** yield the undesired epoxide **3b**. Even though, the initial activity of the whole-cell biocatalyst with the transmembrane enzyme PpGPo1AlkB towards isoprenyl acetate **2** was rather low (0.28 U g_CDW_ ^-1^), we were able to achieve a 6-fold improvement by applying rational design and transporter co-expression. Notably, the exchange I233V in PpGPo1 (I238V in M_AlkB) seems to boost the catalytic efficiency independently of the substrate in both homologsThis mutation was identified by Koch *et al*. after two rounds of directed evolution of PpGpo1AlkB in a triple variant (V129M+L132V+I233V (MVV)) showing higher activities towards short-chain alkanes (C_4_-C_5_) than the L132V single mutant found in the first round of evolution.^[37]^ Generally, we observed that the amino acid substitutions have a similar effect in the different AlkB homologs, and I238V and F169L led to a comparable increase in activity (**Table 1**). A similar transferability within this enzyme family was shown for the W55 residue (PpGPo1AlkB). In several AlkB monooxygenases, the exchange to smaller amino acids such as serine or cysteine at this position shifted the substrate acceptance towards longer alkanes.^[28,36]^

In the docking studies the position F164 in PpGPo1 was identified to be in close proximity to the ligand **2** and exhibiting high REU. As shown in **Figure 3**, the exchange to Leu led to increased activity towards **2** but a decrease with *n*-octane. The exchange to isoleucine or valine led to lower activities with both substrates and the formation of not only **3a** but also a closely, not yet identified related side-product **3x** (**Figure S7**). Interestingly, **3x** is also formed by the M_AlkB variants F169L/I/V. Further detailed analyses are necessary for a clearer understanding of the underlying mechanism of these enzymes. As a general trend, we could observe a shift in substrate preference *P*_Var_⁄*P*_WT_ to isoprenyl acetate **2** over *n-*octane in the modelling-derived variants (**Figure 3**). This effect was most profound in the A268F variant (*P*_Var_⁄*P* = 15.1; **Table S8**). The absolute activity towards **2** was in fact lower than WT-activity but the activity towards *n-*octane dropped to 0.08 U g_CDW_ ^-1^.

Tulipalin A offers highly promising features as a biobased monomer and as a potential substitute for petrol-based methacrylic acid for several applications. The proposed downstream pathway links its biosynthesis to isoprenyl pyrophosphate, a readily available metabolite (**Scheme 1**). The key for this pathway is the herein demonstrated regioselective hydroxylation of isoprenyl acetate **2** at the C4-position without undesired oxidation of the *exo*-olefin group. The bacterial alkane monooxygenase AlkB displays excellent selectivity for terminal hydroxylation, thus unlocking the new biosynthetic pathway. Furthermore, we demonstrate that the activity of membrane-bound monooxygenase towards the sterically demanding substrate can be improved by rational enzyme engineering. Combined with transport engineering, this strategy significantly improved the whole-cell biocatalyst, unlocking an enzymatic pathway for the fermentative production of the biobased tulipalin A.

## Experimental

The conversion of **2** to 4-acetoxy-2-methylene-butan-1-ol **3a** was performed as whole-cell biotransformation. *E. coli* BL21(DE3) cells were transformed with pa*lkB(homolog/mutant)FGT(L)* based on the pCom10 vector backbone which exploits the natural alkane inducible *alkB-* promotor.^[34]^ The main culture was grown in M9 minimal medium, containing 2-fold amounts of NH_4_Cl at 30 °C to an OD_600_ of 0.4 - 0.5 and then induced with dicyclopropyl ketone (0.05 % (v/v)). After 4 h of expression at 30 °C the cells were harvested and resuspended in resting cell buffer (50 mM KPi, pH 7.4, 2 mM MgSO_4_, 1 % glucose) to an OD_600_ ≈ 10. The reaction was initiated by the addition of substrate (5 mM) and performed at 25 °C, 180 rpm at a total volume of 300 µL (for each sampling point). Samples (250 µL) were quenched by adding HCl (2 M, 25 µL) and then extracted with ethyl acetate (250 µL) containing methyl benzoate as internal standard (1 mM). The samples were subjected to GC-MS and/ or GC-FID analysis. Due to the lack of an authentic standard for **3a**, NMR-spectroscopy was done to verify the identity of the product peak appearing in the GC analysis (**Figure S5**). Therefore, the reaction was scaled up to 20 mL in shake flasks and the product was extracted after 24 h of reaction and purified via column chromatography. The recorded spectra was in agreement with reported data.^[35]^ Enzymatic activities (U g_cdw_ ^-1^) were obtained from biological triplicates and calculated as µmol of product formed per min and g_cdw_. The data is represented as arithmetic mean and standard deviation.

## Supporting Information

The authors have cited additional references within the Supporting Information.^[45-53]^

## Acknowledgements

The authors would like to thank Prof. Dr. Anton Glieder (Graz University of Technology and bisy GmbH) and Dr. Kai Baldenius (Baldenius Biotech Consulting) for interesting discussions on unspecific peroxidases. We thank the COMET center acib: Next Generation Bioproduction is funded by BMK, BMDW, SFG, Standortagentur Tirol, Government of Lower Austria, and Vienna Business Agency in the framework of COMET - Competence Centers for Excellent Technologies. The COMET-Funding Program is managed by the Austrian Research Promotion Agency FFG. VD and GO were supported by funding from the European Research Council through a Starting Grant (HelixMold 802217).

## Notes

Supporting information for this article is given via a link at the end of the document.

### Competing Interest Statement

The authors have declared no competing interest.

