## Supplemental Information for "Engineering of Transmembrane Alkane Monooxygenases to Improve a Key Reaction Step in the Synthesis of Polymer Precursor Tulipalin A"

[d] BioTechMed-Graz, Mozartgasse 12/II, 8010 Graz, Austria.

[e] Henkel AG & Co. KGaA, Henkel-Str. 67, 40589 Düsseldorf, Germany

Supporting information for this article is given via a link at the end of the document.

### 1 Experimental information

#### 1.1 Computational methods

The workflow of the docking and computational mutagenesis study is shown in **Figure S1**. The docking study began by preparing the protein scaffolds and ligands (isoprenyl-acetate, and *n*-octane). Protein scaffolds for PpGPo1AlkB and M_AlkB were constructed using AlphaFold v2.3.1 (AF2), ensuring 50 %-70 % sequence coverage with pLDDT scores > 90 %. Using Rosetta the iron atoms were added to the protein scaffold by placing two additional virtual atoms to the iron ligand params files to ensure constraint geometries were measured accurately.[45] This parameter file was used for matching to the AF2 predicted structure. Metal atom placement was finalized using RosettaScripts employing the AddOrRemoveMatchCsts and EnzRepackMinimize movers to ensure correct coordination to the histidine residues—138 and 273 for PpGPo1AlkB and 143 and 278 for M_AlkB.

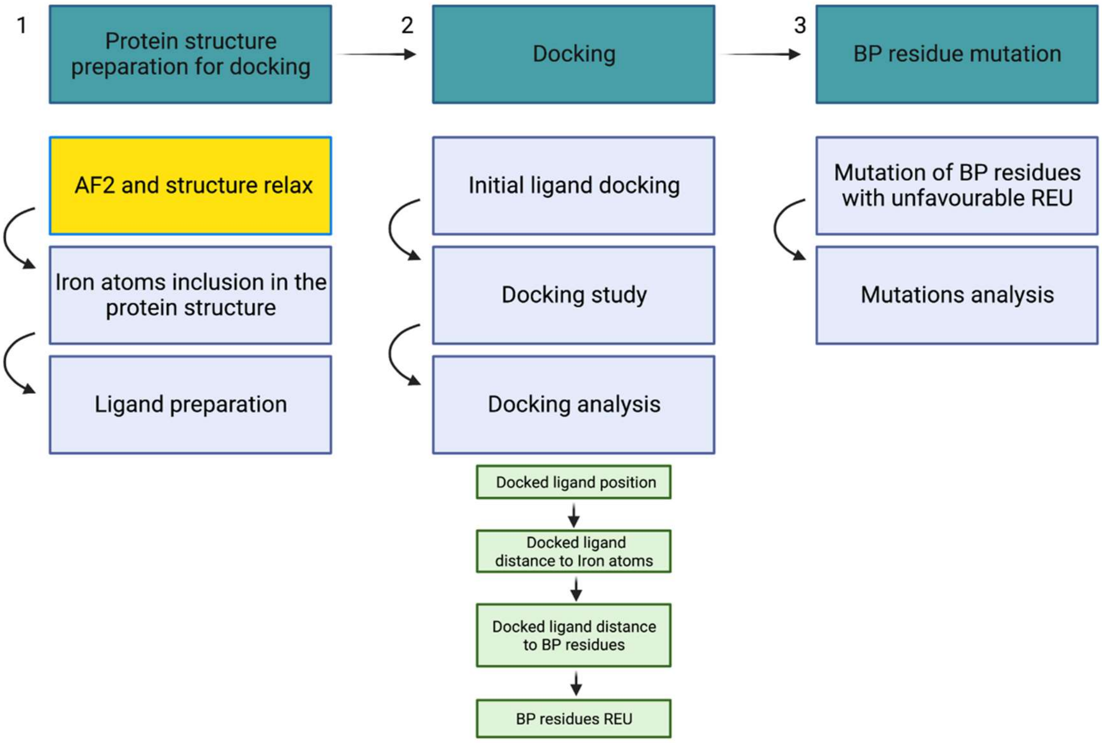

**Figure S1:** Workflow of computational docking and mutagenesis study of AlkB.

Parameter and coordinate files of the ligand molecules were downloaded from PubChem in XML and PDB formats. The files were converted to mol2 format using OpenBabel[46] followed by conversion to Rosetta params files using the molefile_to_params.py script, as provided with the Rosetta modeling suite.

Initial ligand placement was performed via geometry-based molecular docking using the PatchDock server, with the best-scored ligand pose selected for further high-resolution docking. High-resolution docking was restricted to the central pocket, near the iron atoms, due to constraints such as the protein models' rigidity and lack of favorable docking regions elsewhere, aiming to identify important residues for enhancing catalytic activity. RosettaScripts was used to assemble a full docking protocol, which included initial minimization and repacking of the protein scaffold, followed by the execution of the GALigandDock[47] mover for docking, and concluded with a final round of structure minimization and repacking. Data analysis filtered out ligands docked outside the protein scaffold. Evaluation of docked ligands relied on the Rosetta Ligand score and geometries of the docked structure. The total score of the best-scoring molecules was used for further analysis. Approximately 90% of the docked ligands retained final positions within the protein scaffold. This enabled the identification of potential key residues for enzymatic activity by docking of isoprenyl acetate and *n*-octane, respectively. These residues were identified via their distances between the terminal atoms in the docked ligand poses and the surrounding residues. For PpGPo1AlkB, surrounding residues included TRP 55, ILE 128, LEU 132, ASN 135, PHE 164, GLN 264, LEU 265, ALA 268, ASN 269, ILE 271, VAL 305, LEU 306, LEU 309, TYR 339, and PHE 343. For M_AlkB, they comprised GLY 136, LEU 139, PHE 166, TYR 170, GLU 172, GLU 203, ILE 204, ALA 207, PHE 208, TRP 267, LEU 270, THR 271, and ASN 274. The nearest residues to the ligands' terminal carbon atoms were identified to be ASN 135, PHE 164, LEU 265, ALA 268, and LEU 309 for PpGPo1AlkB and GLY 136, LEU 139, TYR 169, ILE 204, LEU 270, and ASN 274 for M_AlkB. Consistency was ensured by calculating average distances between these residues and ligand atoms of the best-scored ligands. Additionally, we assessed the Rosetta score per residue of these closest residues. For PpGPo1AlkB, ASN 135, LEU 265, LEU 309, and TYR 339 exhibited positive scores, indicating unfavorable energetic conditions for ligand docking. For M_AlkB, GLY 136 and ILE 204 showed similar unfavorable conditions, guiding the selection process for key residues involved in catalysis.

The selected residues were optimized via RosettaScripts utilizing the FastDesign mover, which selectively redesigns them using a limited amino acid alphabet. Output sequences are compared to evaluate the frequency of occurrence for each substituent, indicating structural favorability. For PpGPo1AlkB, the analysis reveals LEU as the most frequent residue at positions 135, 164, and 309, TRP at position 265, and ILE at position 268, while the native amino acid remains predominant at position 339. For M_AlkB, the most frequently occurring substituents are MET at position 136, LEU at 139, PHE at 169, LEU at 204, PHE at 270, and LEU at 274.

Full code is available under: https://github.com/veronicadelsoglio-code/Docking-study-AlkB/blob/main/Mutation.xml.

#### 1.2 Chemicals and devices

If not stated otherwise, all chemicals and compounds used in this work were either purchased from Merck/Sigma-Aldrich, TCI chemicals or Carl Roth usually in the highest purity available. Standard laboratory equipment has been used for this study.

##### 1.2.1 Synthesis of isoprenyl acetate 2

**2** was synthesized from isoprenol **1** similarly as described by Bartoli and co-workers.[48] To a 50 mL round-bottom flask equipped with magnetic stirring, 60.9 mmol of $Ac_2O$ and 0.58 mmol of $Mg(ClO)_4$ were added. The reaction was initiated by dropwise addition of **1** (58 mmol). The reaction was monitored by TLC (cyclohexane/EtOAc 3:1) until completion (approx. 30 min). Aqueous $NaHCO_3$ was added, and the product was extracted with $Et_2O$. The organic layer was dried over $MgSO_4$ and the solvent was evaporated to give the pure acetate ester (yield: 88%). The formation of **2** was verified by GC-MS and NMR-spectroscopy and compared to reported spectroscopic data ($^1$H NMR (300 MHz, $CDCl_3$), *δ* 4.81 (s, 1H), 4.74 (s, 1H), 4.18 (t, *J* = 6.87 Hz, 2H), 2.34 (t, *J* = 6.72 Hz, 2H), 2.04 (s, 3H), 1.76 (s, 3H) ppm).[49]

##### 1.2.2 Synthesis of 4-acetoxy-2-methylene butyric acid 5

In a 10 mL round bottom flask equipped with a magnetic stirrer 200 mg (1.7 mmol) of 2-methylen-4-hydroxy butyric acid (Enamine, Kyiv, Ukraine) were selectively acetylated to 4-acetoxy-2-methylene butyric acid **5** with acetic anhydride (5.0 eq.) and pyridine (5.7 eq.) at 0 °C overnight. After completion of the reaction (monitored by TLC), the reagents were initially evaporated *in vacuo*. The obtained residue was washed with 1 M aq. HCl (5 mL) and extracted with EtOAc (3 x 5 mL). The combined organic layers were concentrated under reduced pressure in a water bath at 50 °C. Further evaporation under inert gas conditions to remove remaining traces of reagents yielded 4-acetoxy-2-methylene butyric acid **5** in pure form as a yellow, viscous oil (167 mg, 62% yield). NMR data of the product is in total agreement with the literature ($^1$H NMR (300 MHz, $CDCl_3$), *δ* 6.37 (s, 1H), 5.73 (s, 1H), 4.23 (t, *J* = 6.6 Hz, 2H), 2.65 (t, *J* = 6.6 Hz, 2H), 2.03 (s, 3H) ppm).[50]

##### 1.2.3 Synthesis of 3-methyl-3,4-epoxybutyl acetate 3b

2-(2-Methyl-2-oxiranyl)ethanol **2b** (Enamine, Kyiv, Ukraine) was acetylated using a lipase as catalyst to the predicted product 3-methyl-3,4-epoxybutyl acetate **3b** using vinyl acetate. Briefly, 20.4 mg (0.20 mmol) of **2b** was reacted with 37 µL (0.40 mmol) of vinyl acetate in dry MTBE following previously published methods.[35,51] 1.62 mg Lipase from *Pseudomonas cepacia*, >30 U mg$^{-1}$, Sigma-Aldrich) was added and the reaction mixture was performed at 30 °C and 500 rpm for 24 h. The reaction mixture was centrifuged, and the supernatant was concentrated using a rotary evaporator. Analysis of the reaction mixture was performed using GC-MS and further $^1$H NMR $CDCl_3$, 300 MHz: δ 1.29, (s, 3H), δ1.74-1.94 (m, 2H), *δ*1.99 (s, 3H), *δ*2.53 (d, 1H, *J* = 6 Hz), δ2.58 (d, 1H, *J* = 4.6), *δ* 4.00-4.19 (m, 2H).

#### 1.3 Cloning of *alkB*-operon

Synthetic DNA-fragments encoding *alkBFG*, *alkT, alkJ* and *alkH* from *P. putida* GPo1 were ordered from Integrated DNA Technologies (IDT; Leuven, Belgium). The sequence information was taken from the *P. putida* OCT plasmid *alk* genes cluster (NCBI nucleotide: AJ245436.1). The fragments were sub-cloned into the pJET1.2 vector using the Thermo Scientific CloneJET PCR Cloning Kit. The *alkBFG* and *alkT* fragments were inserted into the pCom10 vector backbone via the FastCloning method.[34,52] The operon of the p*alkBFGT* vector was expanded with *alkL, alkHJ, alkHJL* and *alkJ,* resulting in p*alkBFGTL, palkBFGTHJ, alkBFGTHJL, and palkBFGTJ,* respectively. The *alkL* gene was amplified from the pCom10_AlkL[34] adding an RBS upstream of *alkL* and subsequently cloned into the p*alkBFGT* via FastCloning. The constructs *palkBFGTHJ, alkBFGTHJL, and palkBFGTJ* were generated via Gibson Assembly by expanding the operon with the respective fragments as reported by Nuland *et al*.[27,53]

Four additional genes encoding for homologous AlkBs from other organisms were also ordered as synthetic DNA-fragments from IDT: a) AlkB from *Pseudomonas putida* P1 (PpP1AlkB; NCBI protein: CAB51047.1)[39] b) AlkB1 from *Alcanivorax borkumensis* (AboAlkB; NCBI protein: BAC98365.1)[40] and c) AlkB from *Acinetobacter baylyi* (AboAlkB; NCBI protein: WP_120429654.1) and d) M_AlkB from *Marinobacter* sp. (M_AlkB; NCBI protein: MAB50652.1). The genes of the homologous *alkBs* were cloned into the p*alkBFGT* vector via Gibson Assembly replacing the *PpGpo1alkB*, while leaving the electron transfer system *PpGPo1alkFGT* intact. Correct sassembly was verified by Sanger Sequencing (Microsynth, Vienna). **Table S1** lists all plasmids used in this study.

**Table S1:** Listing of plasmids used in this work. All plasmids for the expression of *alkB(homolog/mutant)* are based on the broad-host vector pCom10 employing an alkane-inducible promotor system.[34] The inserts have been ordered as synthetic genes and cloned into the pCom10-backbone either by the FastCloning method or Gibson Assembly.

| Name | Gene origin | Backbone | Purpose |
|---|---|---|---|
| pPpGpo1alkB-FGT | *P. putida* GPo1 | pCom10 | Expression of PpGpo1*alkBFGT* |
| pPpGpo1alkB-FGTL | *P. putida* GPo1 | pCom10 | Expression of PpGpo1*alkBFGTL* |
| pPpGpo1alkB-FGTHJ | *P. putida* GPo1 | pCom10 | Expression of PpGpo1*alkBFGTHJ* |
| PpGpo1alkBF-GTHJL | *P. putida* GPo1 | pCom10 | Expression of PpGpo1*alkBFGTHJL* |
| pPpGpo1alkB-FGTJ | *P. putida* GPo1 | pCom10 | Expression of PpGpo1*alkBFGTJ* |
| pPpP1AlkB-FGT | *P. putida* P1 | pPpGpo1alkB-FGT | Expression of *PpP1alkB*-PpGpo1*alkFGT* |
| pM_AlkB-FGT | *Marinobacter sp.* | pPpGpo1alkB-FGT | Expression of *M_alkB*-PpGpo1*alkFGT* |
| pAboAlkB-FGT | *A. borkumensis* | pPpGpo1alkB-FGT | Expression of *AboalkB*-PpGpo1*alkFGT* |
| pAbaAlkB-FGT | *A. baylyi* | pPpGpo1alkB-FGT | Expression of *AbaalkM*-PpGpo1*alkFGT* |
| pPpP1AlkB-FGTL | *P. putida* P1 | pPpGpo1alkB-FGTL | Expression of *PpP1alkB*-PpGpo1*alkFGTL* |
| pM_AlkB-FGTL | *Marinobacter sp.* | pPpGpo1alkB-FGTL | Expression of *M_alkB*-PpGpo1*alkFGTL* |
| pAboAlkB-FGTL | *A. borkumensis* | pPpGpo1alkB-FGT | Expression of *AboalkB*-PpGpo1*alkFGTL* |
| pAbaAlkB-FGTL | *A. baylyi* | pPpGpo1alkB-FGT | Expression of *AbaalkM*-PpGpo1*alkFGTL* |
| pCom10_AlkL | *P. Putida* GPo1 | pCom10 | Backbone for cloning |
| pCom10_empty | - | pCom10 | Empty vector control (EVC) |
| pJET1.2_alkBFG | *P. putida* GPo1 | pJET1.2 | Cloning of synthetic DNA fragments |
| pJET1.2_alkT | *P. putida* GPo1 | pJET1.2 | Cloning of synthetic DNA fragments |
| pJET1.2_AlkH | *P. putida* GPo1 | pJET1.2 | Cloning of synthetic DNA fragments |
| pJET1.2_AlkJ | *P. putida* GPo1 | pJET1.2 | Cloning of synthetic DNA fragments |

Single and combinatorial mutants of PpGPo1AlkB and M_AlkB (**Table S2**) were generated by site-directed mutagenesis (SDM). The non-active variant H273A was generated for control experiments. Successful mutagenesis was verified by Sanger sequencing.

**Table S2:** Listing of all mutants of PpGpo1AlkB and M_AlkB tested in this work. Mutants were created by SDM using pPpGpo1alkB-FGT(L) or pM_alkB-FGT(L) as template.

| Variants PpGpo1AlkB | Ref. | Variants PpGpo1AlkB | Ref. |
|---|---|---|---|
| W55S | [36] | F164L* | this study |
| V129M | [37] | F164W | this study |
| L132V | [37] | F164I | this study |
| I233V* | [37] | F164V | this study |
| H273A | [36] | L265W | this study |
| V129M + L132V + I233V (MVV) | [37] | L265Y | this study |
| W55S + V129M + L132V + I233V (MVV) | [36,37] | A268I | this study |
| N135L | this study | A268F | this study |
| N135F | this study | F164L + I233V (LV)* | this study |
| **Variants M_AlkB** | **Ref.** | **Variants M_AlkB** | **Ref.** |
| I238V* | this study | F169L* | this study |
| F169L + I238V (LV)* | this study | | |

* mutant also in pPpGpo1AlkB-FGTL/ pM_AlkB-FGTL

### 1.4 UPO biotransformations

The enzyme panel consisting of 77 UPOs was purchased from Aminoverse (Nuth, The Netherlands). The kit contains 62 wildtype and 15 mutant fungal LPOs (**Table S3**) which were recombinantly produced as secreted protein by *Pichia pastoris* and were available in lyophilized form (1 mg) stored at -20 °C before use. Isoprenyl acetate (100 mM) was prepared in acetonitrile and the co-substrate $H_2O_2$ (55 mM) in 1x tricine buffer (100 mM, pH 7.5). The reaction mixture containing 195 µL buffer, resuspended UPO solution (25 µL), and 10 µL isoprenyl acetate stock solution, was placed in a reaction tube. The reaction was initiated by the addition of 5 µL $H_2O_2$ stock solution. The reaction vial was placed in a Thermoshaker operated at 30 °C and 500 rpm. 5 µL $H_2O_2$ was pulsed every 30 minutes until 2 h reaching a concentration of 4.4 mM. The reaction was carried out until 4 h and quenched with the addition of the UPO-STOP solution (2 µL) containing catalase and further incubated at room temperature for at least 15 min. The reaction mixture was then extracted with ethyl acetate containing 1 mM of methyl benzoate as an internal standard for GC-MS measurements.

A large-scale reaction (155x) was set up to confirm the structure of the UPO-product by NMR-spectroscopy. Conversion of **2** by UPO 12 Aminoverse/ UPO 13 Bisy (provided by Prof. Dr. Anton Glieder) was performed for 24 h at 30 °C using 1.1 equivalence of $H_2O_2$. The product was extracted and purified from the reaction mix. After 24 h, the reaction was quenched with the addition of catalase (43 µL, 20,000 U $mL^{-1}$). The product was then extracted using dichloromethane (1:2 v/v, 6x) and subjected to GC-MS and $^1$H NMR analysis.

**Table S3:** Plate layout of UPO Enzyme panel (Aminoverse (Nuth, The Netherlands)).

| | | | | | | | | | |
|---|---|---|---|---|---|---|---|---|---|
| UPO 1 | UPO 5 | UPO 3 | UPO 21 | UPO 29 | UPO 42 | ***Aae* UPO*** | UPO 56 | UPO 13M2 | UPO 36M1 |
| UPO 4 | UPO 8 | UPO 28 | UPO 24 | UPO 32 | UPO 38 | UPO 49 | UPO 57 | UPO 13M5 | UPO 36M2 |
| UPO 7 | UPO 11 | UPO 31 | UPO 19 | UPO 26 | UPO 44 | UPO 50 | UPO 58 | UPO 13M4 | UPO 36M3 |
| UPO 10 | UPO 14 | UPO 6 | UPO 34 | UPO 45 | UPO 27 | UPO 51 | UPO 59 | UPO 13M7 | UPO 36M4 |
| UPO 13 | UPO 17 | UPO 9 | UPO 37 | UPO 46 | UPO 48 | UPO 52 | UPO 60 | UPO 13M8 | UPO 36M5 |
| UPO 16 | UPO 20 | UPO 12 | UPO 39 | UPO 35 | UPO 30 | UPO 53 | UPO 61 | UPO 13M9 | neg. c. ** |
| UPO 22 | UPO 25 | UPO 15 | UPO 41 | UPO 47 | UPO 33 | UPO 54 | UPO 13M3 | UPO 13M10 | UPO 13 |
| UPO 2 | UPO 23 | UPO 18 | UPO 43 | UPO 40 | UPO 36 | UPO 55 | UPO 13M1 | UPO 12M11 | - |

* AaeUPO- from *Agrocybe aegerite*; ** negative control without UPO; M … UPO-mutant

### 1.5 Recombinant expression of *AlkBFGT(L)*

*E. coli* BL21(DE3) harboring the respective plasmid were used for expression. The conditions were set similarly as described in the literature.[27,31] *E. coli* BL21(DE3) pCom10_*empty* was used for negative control experiments. Strains were grown in either LB or M9 minimal medium (**Table S4**) supplemented with 50 µg $mL^{-1}$ kanamycin for selection. 5 mL of LB medium was inoculated with a single colony and incubated overnight at 30 °C and constant agitation (120 rpm). 200 µL of the overnight culture (ONC) were transferred to 20 mL of M9 minimal in a 100 mL baffled shake flask and the culture was again incubated overnight at 30 °C. For the expression, 400 mL of M9 minimal medium were inoculated to an $OD_{600}$ of 0.15 with the M9-preculture and grown at 30 °C until an $OD_{600}$ of 0.4 to 0.5. By adding 0.05 % (v/v) dicyclopropyl ketone (DCPK) recombinant gene expression was induced. After 4 h of expression at 30 °C the cells were harvested by centrifugation (4,400 x *g*, 4 °C, 15 min). The production of the recombinant proteins was verified by SDS-PAGE.

**Table S4**: Compostion of M9 minimal medium. For 1 L of M9 medium, 200 mL of 5x M9-salts, 2 mL of 1M $MgSO_4$, 1 mL of thiamine, 50 µL of biotin, 1 mL of $US^{Fe}$ trace element solution, and 25 mL of glucose solution were mixed and filled up to 1 L with autoclaved $ddH_2O$.

| Compound | Final conc. | Stock | Stock conc. | Stock preparation |
|---|---|---|---|---|
| $Na_2HPO_4$*2 $H_2O$ | 8.5 g $L^{-1}$ | 5x | 42.5 g $L^{-1}$ | pH adjusted to 7.0 with 5 M NaOH; autoclaved; stored at room temperature |
| $KH_2PO_4$ | 3.0 g $L^{-1}$ | | 15 g $L^{-1}$ | |
| NaCl | 0.5 g $L^{-1}$ | | 2.5 g $L^{-1}$ | |
| $NH_4Cl$ | 2.0 g $L^{-1}$ | | 10.0 g $L^{-1}$ | |
| $MgSO_4$*7 $H_2O$ | 2 mM | 1 M | 246 g $L^{-1}$ | Autoclaved, stored at room temperature |
| Thiamine*HCl | 1 mg $L^{-1}$ | 1000x | 1 mg $mL^{-1}$ | 50 mg dissolved in 50 mL $ddH_2O$ and filter sterilized; stored -20°C |
| Biotin | 5µg $L^{-1}$ | 20000x | 0.1 mg/mL | 5 mg dissolved in 45 mL $ddH_2O$, 1 N NaOH added until dissolved, filled up to 50 mL with $ddH_2O$ and filter sterilized, stored at -20 °C |
| $US^{Fe}$ trace element solution* | 1 mL $L^{-1}$ | 1000x | | All components below dissolved in 1 M HCl; filter sterilized; stored at -20°C 1 mL |
| Glucose | 5 g $L^{-1}$ | 20 % | 200 g $L^{-1}$ | Dissolved in 1 L $ddH_2O$; autoclaved; stored at 4 °C |
| ***1000x USFe trace element solution** | | | **Stock conc.** | |
| $FeSO_4$*7 $H_2O$ | | | 8.87 g $L^{-1}$ | |
| $CaCl_2$*2 $H_2O$ | | | 4.12 g $L^{-1}$ | |
| $MnCl_2$*2 $H_2O$ | | | 1.23 g $L^{-1}$ | |
| $ZnSO_4$*7 $H_2O$ | | | 1.87 g $L^{-1}$ | |
| $H_3BO_3$ | | | 0.30 g $L^{-1}$ | |
| $Na_2MoO_4$*2 $H_2O$ | | | 0.25 g $L^{-1}$ | |
| $CuCl_2$*2 $H_2O$ | | | 0.15 g $L^{-1}$ | |
| Disodium EDTA*2 $H_2O$ | | | 0.84 g $L^{-1}$ | |

#### 1.6 AlkB whole-cell biotransformations

After harvest the cells were resuspended in resting cell buffer (RCB; 50 mM KPi, pH 7.4, 1 % glucose, 2 mM $MgSO_4$) if not stated otherwise to an $OD_{600}$ of 10. The reaction mixture was prepared on ice as a homogenous master mix for each time-lapse sample. The reaction was initiated by the addition of 5 mM of substrate (isoprenyl acetate or *n*-octane; 200 mM stock in EtOH; 2.5 % (v/v) EtOH in the reaction) and then separated to 300 µL in 1.5 mL tightly sealed glass vials, which were placed lying in a specialized rack and incubated at 25 °C under constant agitation (180 rpm). 250 µL were sampled (usually after 0, 15, 30, 60, 180 min and 24 h) and quenched by adding 25 µL of 2 M HCl. The samples were stored at -20 °C until further use. The biotransformations were performed in biological triplicates. The data is represented as arithmetic mean and standard deviation. *E. coli* BL21(DE3*)* pCom10_*empty* served as negative control.

For identification of the product formed by AlkB when using isoprenyl acetate as substrate, the reaction was scaled up to a total volume of 20 mL in an Erlenmeyer flask sealed with a metal cap and Parafilm®. The reaction was initiated by adding 5 mM of **2** and performed at 30 °C, 180 rpm for 48 h. The reaction was followed by GC-MS analysis. The product was extracted with EtoAc from the reaction mixture. The organic layer was dried over $MgSO_4$ and the solvent evaporated. The crude was purified with column chromatography (cyclohexane/EtOAc 4:1) to yield the purified compound **2** (Yield: < 5 %; TLC: cyclohexane/EtOAc (2:1)). $^1$H NMR was recorded in $CDCl_3$.

#### 1.7 Lactonization of 4-acetoxy-2-methylene-butanoic acid to tulipalin A

10 mg of **5** were dissolved $ddH_2O$ acidified with HCl (10 eq.) in a total volume of 1 mL. The reaction mix was incubated at 80 °C and 800 rpm in a thermoshaker. 20 µL samples were taken after at the beginning, after 3 h and 20 h and analyzed with TLC (EtoAc/ DCM (3:1) and GC-FID. The samples were extracted with 100 µL EtoAc (containing 1 mM methyl benzoate as internal standard (ISTD)) and further diluted with solvent to 200 µL. To prevent spontaneous lactonization of **5** to **6** during GC-analysis the sample was derivatized by silylation using N,O-bis(trimethylsilyl)trifluoroacetamide (BSTFA).

#### 1.8 GC-analysis

For the analysis of whole-cell biotransformation, 250 µL of the samples were extracted with EtoAc containing methyl benzoate as ISTD (1:1, v/v) by vigorous shaking for 1 min. Phase separation was achieved by centrifugation (16,000 x *g*, 4 °C, 7 min). The organic phase was dried over $NaSO_4$. 200 µL of extract were directly subjected to GC-analysis. Samples containing **5** were derivatized with BSTFA prior to analyis. Therefore, 2 µL of BSTFA were added to the extracted sample and incubated for 1 h at 60 °C. GC−MS analysis was used to qualitatively confirm the formation of the desired products. Conditions for analytics were established using commercially available compounds (**Table S5**). The measurements were performed on a Shimadzu GCMS-QP2010 SE instrument equipped with an AOC-20i/s autosampler and injector unit together with a Zebron ZB-5MSi capillary column (30 m × 0.25 mm x 0.25 µm, Phenomenex).

**Table S5:** Parameters of GC-MS methods (long and short) and retention times of compounds used.

| GC parameters | Method I (short) | Method II (long) |
|---|---|---|
| Column | Zebron ZB-5MSi (Phenomenex, 30 m x 0.25 mm x 0.25 µm) | |
| Flow Control Mode | Linear velocity (39.5 cm $sec^{-1}$) | |
| Total flow/ Column flow/ Carrier gas | 15 mL $min^{-1}$/ 1.21 mL $min^{-1}$/ Helium | |
| Injection temperature/ injection volume/ split ratio | 250 °C/ 1 µL/ 9.1 | |
| Temperature program | 3 min 50 °C, 30 °C $min^{-1}$ to 300 °C, 3 min 300 °C | 5 min 50 °C, 40 °C $min^{-1}$ to 300 °C, 5 min 300 °C |
| **Total program time** | 14.33 min | 16.25 min |
| **MS parameters** | **Method I (short)** | **Method II (long)** |
| Ion source temperature | 250 °C | |
| Interface temperature | 320 °C | |
| Mode | Scan, 30 – 300 m $z^{-1}$ | |
| **Compounds** | **Rt Method II (long) [min]** | **Rt Method I (short) [min]** |
| Methyl benzoate (ISTD) | n. d. | 6.55 |
| Dicyclopropyl ketone (DCPK; inducer) | 6.97 | 5.48 |
| Isoprenyl acetate **2** | 6.15 | 4.77 |
| 4-acetoxy-2-methylene-butan-1-ol **3a** | 8.14 | 6.71 |
| 3-methyl-3,4-epoxybutyl acetate **3b** | n. d. | 6.07 |
| 4-acetoxy-2-methylene-butanal **4** | n. a. | n. a. |
| 4-acetoxy-2-methylene-butyric acid **5** | 8.87 | 7.35 |
| Silylated 4-acetoxy-2-methylene-butyric acid **5-TMS** | n. d. | n. d.. |
| Tulipalin A **6** | 7.15 | 6.05 |

n. d.: not determined; n. a.: no authentic standard available

For quantitative analysis the quenched reaction samples were extracted with EtoAc containing 1 mM of ISTD and then subjected to GC-FID analysis on a Shimadzu Nexis GC-2030 equipped with an AOC-20i Plus autosampler and injector unit and a Zebron ZB-5MSi capillary column (30 m × 0.25 mm x 0.25 µm, Phenomenex). The analytical conditions were established using pure compounds (**Table S6**). The concentrations of the analytes were calculated by external calibration curves generated by measuring samples with known concentrations of the pure compounds (0 – 6 mM) extracted from resting cell buffer. Concentrations of compounds that are not available were calculated based on calibration curves of structurally similar chemicals.

**Table S6:** Parameters of applied GC-FID method, retention times and calibration variables of used compounds.

| Parameters | Method isoprenyl acetate | Method *n*-octane |
|---|---|---|
| **Column** | Zebron ZB-5MSi (Phenomenex, 30 m x 0.25 mm x 0.25 µm) | |
| **Flow Control Mode** | Linear velocity (22 cm sec-1) | |
| **Total flow/ Column flow/ Carrier gas** | 15.9 mL min-1/ 1.18 mL min-1/ Nitrogen | |
| **Injection temperature/ injection volume/ split ratio** | 250 °C/ 1 µL/ 10 | |
| **Temperature program** | 1 min 50 °C, 20 °C min-1 to 250 °C, 2 min 300 °C | 1 min 50 °C, 20 °C min-1 to 250 °C, 2 min 300 °C |
| **FID temperature** | 320 °C | |
| **Total program time** | 13.00 min | 10.50 min |

| Rt method isoprenyl acetate | | Rt method *n*-octane [min] | |
|---|---|---|---|
| Methyl benzoate (ISTD) | 7.06 | Methyl benzoate (ISTD) | 5.62 |
| Dicyclopropyl ketone (DCPK) | 5.85 | Dicyclopropyl ketone (DCPK) | 4.88 |
| Isoprenyl acetate **2** | 5.12 | *n-octane* | 3.94 |
| 4-acetoxy-2-methylene-butan-1-ol **3a** | 7.28 | 1-octanol | 5.41 |
| 3-methyl-3,4-epoxybutyl acetate **3b** | 6.49 | Octanal | 5.07 |
| 4-acetoxy-2-methylene-butanal **4** | n. a. | Octanoic acid | 5.92 |
| 4-acetoxy-2-methylene-butyric acid **5** | 8.14 | | |
| Silylated 4-acetoxy-2-methylene-butyric acid **5-TMS** | 8.44 | | |
| Tulipalin A **6** | 6.05 | | |

#### 1.9 NMR-spectroscopy

$^{1}$H- and $^{13}$C-spectra were recorded using an Avance$^{TM}$ III 300 MHz FT NMR spectrometer. For the analysis 5-10 mg of analyte were dissolved in $CDCl_3$. The spectroscopic data was compared to reported spectra.

### 2 Supporting results

#### 2.1 Acetylation of 2-(2-Methyl-2-oxiranyl)ethanol to 3b

As the formation of the epoxide **3b** was expected to be a potential product in the reaction of isoprenyl acetate **2** with oxygenases an authentic standard was needed. Hence, 3-methyl-3,4-epoxybutyl acetate **3b** was synthesized via lipase-catalyzed acetylation of 2-(2-Methyl-2-oxiranyl)ethanol **2b**. GC-MS analysis of the lipase-catalyzed reaction showed a product peak at 6.07 min (**Figure S2**) and also analyzed via NMR. The recorded spectra agreed with literature and predicted data.[20]

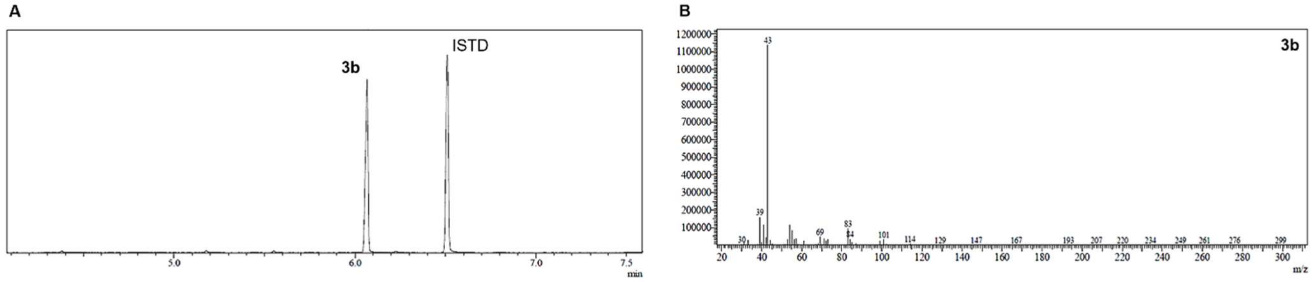

**Figure S2: (A)** GC-MS chromatogram of lipase-catalyzed acetylation of 2-(2-Methyl-2-oxiranyl)ethanol **2b** to 3-methyl-3,4-epoxybutyl acetate **3b** (Rt: 6.07 min) and **(B)** mass spectogram of **3b**.

#### 2.2 UPO-catalyzed transformation of isoprenyl acetate

Unspecific peroxygenases were thought to be interesting candidates to catalyze the terminal hydroxylation of isoprenyl acetate to **3a** – the key reaction in the artificial route towards tulipalin A (**Scheme 1**). In a screening with 77 different UPOs from Aminoverse (Nuth, The Netherlands) we found that 72 UPOs, including *Aae*UPO are active towards **2.** GC-MS analysis (**Figure S3 A-B**) showed that the formed product (Rt: 6.07 min) did not correspond to the desired alcohol **3a** but rather to the epoxide **3b** instead. This was confirmed by comparison with the product from the acetylation of **2b** (**Figure S3 C**). In addition, the NMR of the isolated and purified UPO-product ($^1$H NMR, $CDCl_3$: *δ* 1.29 (s, 3H), *δ* 1.74-1.87 (m, 2H), *δ* 2.34 (s, 3H), *δ* 2.53 (d, 1H, *J* = 4.52), *δ* 2.58 (d, 1H, *J* = 4.56), *δ* 4.01-4.20 (m, 2H)) was in agreement with the NMR of **3b** synthesized by acetylation of **2b**

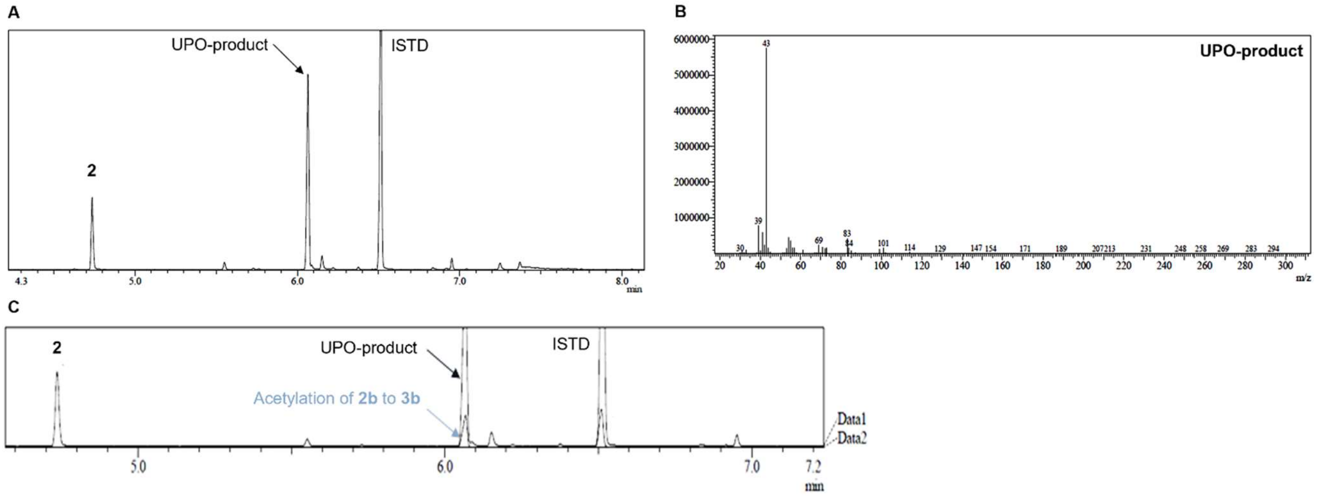

**Figure S3: (A)** GC-MS chromatogram of *Aea*UPO-mediated conversion of isoprenyl acetate **2** gave a prominent product peak at 6.07 min. **(B)** Mass spectogram of the UPO-product. **(C)** Comparison of the chromatogram of the UPO-reaction with **2** and the lipase-catalyzed acetylation of **2b**.

**Table S7** gives a semi-qualitative summary of the formation of **3b** by the 77 UPOs from the enzyme panel. Besides UPO 27, UPO 30, UPO 41, UPO 43 and UPO 48, all other were active towards isoprenyl acetate, with *Aae*UPO yielding the highest amount of **3b**.

**Table S7:** Out of the 77 UPOs (Aminoverse, Nuth, The Netherlands), 72 UPOs converted isoprenyl acetate **2** to the epoxide **3b** but none of the gave the desired terminal alcohol **3a**. +++, ++ and + indicate GC-MS peak areas of **3b** to in the magnitude of ≥ $10^7$, ≥ $10^6$, ≥ $10^5$, while – indicates no product formation.

| | | | | | | | | | |
|---|---|---|---|---|---|---|---|---|---|
| UPO1<br>++ | UPO5<br>+++ | UPO3<br>+++ | UPO21<br>++ | UPO29<br>++ | UPO42<br>++ | *Aae*UPO*<br>+++ | UPO56<br>+++ | UPO13M2<br>+++ | UPO36M1<br>++ |
| UPO4<br>+++ | UPO8<br>+++ | UPO28<br>+ | UPO24<br>+++ | UPO32<br>+++ | UPO38<br>+++ | UPO49<br>+++ | UPO57<br>+++ | UPO13M5<br>+++ | UPO36M2<br>+++ |
| UPO7<br>+ | UPO11<br>+++ | UPO31<br>++ | UPO19<br>+++ | UPO26<br>+ | UPO44<br>++ | UPO50<br>+++ | UPO58<br>+++ | UPO13M4<br>+++ | UPO36M3<br>+++ |
| UPO10<br>++ | UPO14<br>++ | UPO6<br>+ | UPO34<br>++ | UPO45<br>++ | UPO27<br>- | UPO51<br>++ | UPO59<br>++ | UPO13M7<br>++ | UPO36M4<br>++ |
| UPO13<br>++ | UPO17<br>++ | UPO9<br>+ | UPO37<br>++ | UPO46<br>+ | UPO48<br>- | UPO52<br>++ | UPO60<br>++ | UPO13M8<br>++ | UPO36M5<br>++ |
| UPO16<br>++ | UPO20<br>++ | UPO12<br>++ | UPO39<br>+ | UPO35<br>++ | UPO30<br>- | UPO53<br>++ | UPO61<br>++ | UPO13M9<br>++ | neg. control<br>- |
| UPO22<br>++ | UPO25<br>++ | UPO15<br>++ | UPO41<br>- | UPO47<br>++ | UPO33<br>++ | UPO54<br>++ | UPO13M3<br>++ | UPO13M10<br>++ | UPO13<br>++ |
| UPO2<br>++ | UPO23<br>+ | UPO18<br>++ | UPO43<br>- | UPO40<br>+ | UPO36<br>++ | UPO55<br>++ | UPO13M1<br>++ | UPO12M11<br>++ | |

#### 2.3 *alkBFGT(HJL)* expression whole-cell hydroxylation of isoprenyl acetate

Polycistronic expression of the *alk*-operon was confirmed by SDS-PAGE. An example of the expression of *alkBFGTHJL* expression is given in **Figure S4**. AlkB, the outer-membrane transporter AlkL and the membrane-anchored AlkJ were expected in the insoluble fraction. The soluble redox-system AlkFGT and AlkH in the soluble fraction.

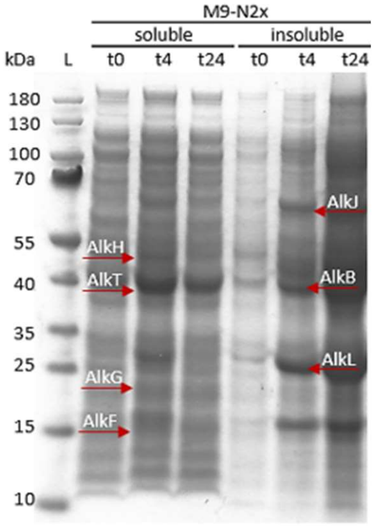

**Figure S4:** SDS-PAGE analysis of the soluble (s) and insoluble fraction of *alkBFGTHJL expression. The membrane (m) proteins are expected in the insoluble fraction.* AlkB: 46 kDa (m), AlkF: 15 kDa (s), AlkG: 19 kDa (s), AlkS: 99 kDa (s), AlkT: 41 kDa (s), AlkL: 25 kDa (m), AlkJ: 61 kDa (m), AlkH: 53 kDa (s).

Whole-cell biotransformations using *E. coli* BL21(DE3) pAlkB(mut/homolog)FGT(L) and **2** and *n*-octane as substrates were qualitatively analyzed by GC-MS and quantitatively by GC-FID. Initial biotransformation with cells expression *PpGPo1alkBFGT and* **2** as substrate showed the formation of an unknown product peak at 6.65 min with a mass spectrogram fitting to the desired terminal alcohol **3a** (**Figure S5**). In order to confirm the formation of 4-acetoxy-2-methylene-butan-1-ol **3a** by AlkB, the product was isolated and purified from whole-cell reactions. The reaction was scaled up to 20 mL (and the product of six reactions was pooled and extracted from with MTBE. The crude extract was subjected to column chromatography to yield pure **2** (less than 5 %). The purification procedure was followed by GC-MS. The formation of **3a** was confirmed via NMR-spectroscopy ($^1$H NMR ((300 MHz, $CDCl_3$), $\delta$ 5.12 (s, 1H), 4.94 (s, 1H), 4.22 (t, J= 6.72 Hz, 2H), 4.11 (s, 2H), 2.42 (t, J= 6.39 Hz, 2H), 2.05 (s, 3H) ppm). The recorded spectrum was in total agreement with the literature.[35]

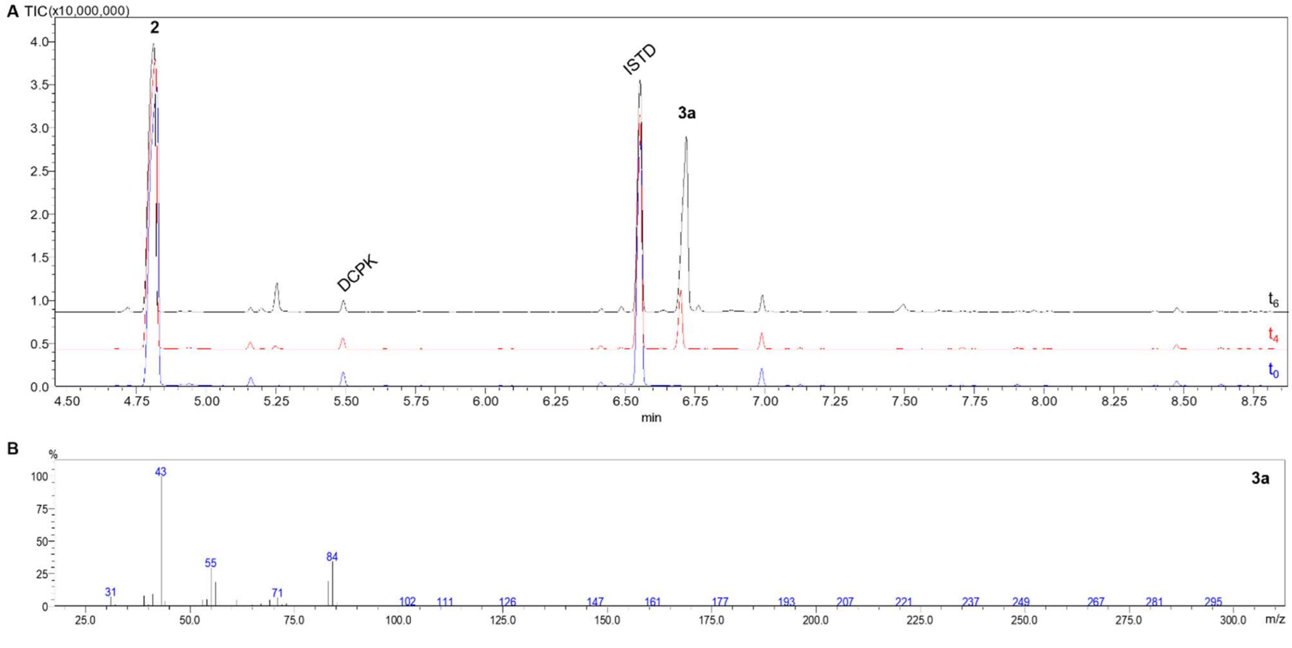

**Figure S5:** (**A**) GC-MS analysis of whole-cell biotransformations with cells expressing PpGPo1AlkB and **2** as substrate (Rt: 4.82 min) shows a peak appearing after 4h at 6.71 min with a mass spectrum (**B**) fitting to the desired product **3a**.

It was assumed that alkane monooxygenases related to AlkB are also able to selectively hydroxylate isoprenyl acetate, some might exhibit even higher activities than AlkB from *P. putida* GPo1. This was exemplified by testing four homologous alkane monooxygenases from *Pseudomonas putida* P1 (PpP1AlkB), *Marinobacter* sp. (M_AlkB), *Alkanivorax borkumensis* (AboAlkB) and *Acetionobacter baylyi* (AbaAlkB). An alignment of the respective protein sequences is shown in **Figure S6**. The different AlkB homologs and variants were tested in whole-cell biotransformations, and the enzymatic activities (U $g_{cdw}^{-1}$) for the two substrates isoprenyl acetate and *n*-octane were determined from biological triplicates and calculated as µmol of product formed per min and $g_{cdw}^{-1}$. The data is represented as arithmetic mean and standard deviation in **Table S8**.

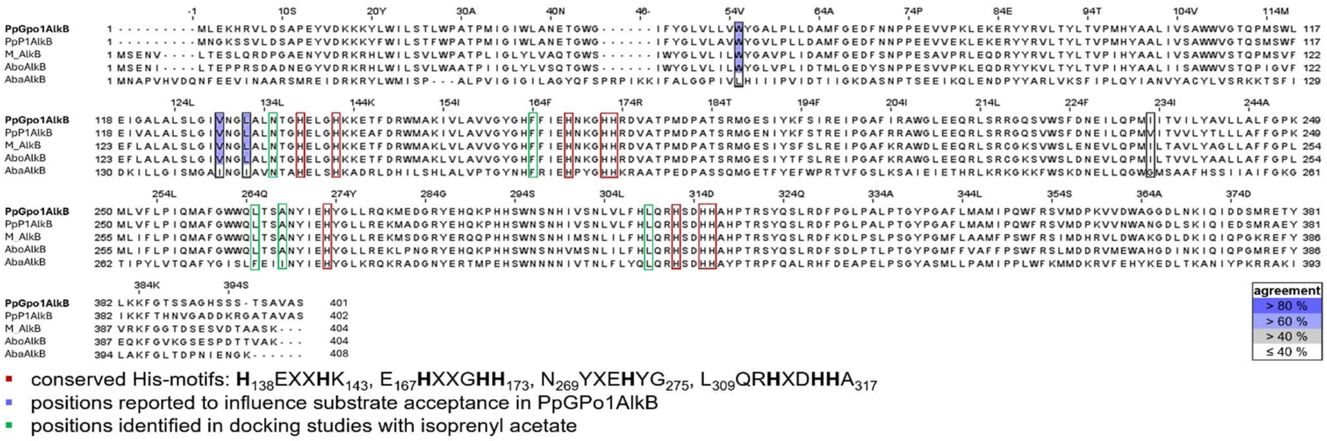

**Figure S6:** Protein sequence alignment of selected AlkB-homologs. The conserved His residues coordinating the iron atoms in the catalytic center are marked with red boxes. Residues that were shown to influence the substrate acceptance in the AlkB from P. putida GPo1 are also highlighted.[36,37]

**Table S8:** Specific activities of AlkB WT and variants without AlkL. The substrate preference $P$ indicates the activity of isoprenyl acetate over *n*-octane, while the preference shift $\frac{P_V}{P_{WT}}$ compares the change of substrate preference of the variant over the wildtype enzyme.

| Enzyme | Variant (V) | Activity [U $g_{cdw}^{-1}$] Isoprenyl acetate | Activity [U $g_{cdw}^{-1}$] *n*-octane | Preference $P$ | Preference shift $P_V/P_{WT}$ |
|---|---|---|---|---|---|
| PpGPo1AlkB | WT | 0.28 ± 0.04 | 4.43 ± 0.16 | 0.1 | 1.0 |
| PpGPo1AlkB | W55S [36] | 0.04 ± 0.00 | 1.11 ± 0.10 | 0.0 | 0.5 |
| PpGPo1AlkB | V129M [37] | 0.00 ± 0.00 | 2.95 ± 0.45 | 0.0 | 0.0 |
| PpGPo1AlkB | L132V [37] | 0.00 ± 0.00 | 1.77 ± 0.54 | 0.0 | 0.0 |
| PpGPo1AlkB | I233V [37] | 0.74 ± 0.06 | 4.83 ± 0.79 | 0.2 | 2.4 |
| PpGPo1AlkB | MVV [37] | 0.00 ± 0.00 | 1.92 ± 0.37 | 0.0 | 0.0 |
| PpGPo1AlkB | SMVV [36,37] | 0.00 ± 0.00 | 0.05 ± 0.04 | 0.0 | 0.0 |
| PpGPo1AlkB | N135L | 0.00 ± 0.00 | 1.11 ± 0.04 | 0.0 | 0.0 |
| PpGPo1AlkB | N135F | 0.00 ± 0.00 | 0.00 ± 0.00 | - | - |
| PpGPo1AlkB | F164L | 0.82 ± 0.06 | 2.81 ± 0.08 | 0.3 | 4.6 |
| PpGPo1AlkB | F164W | 0.38 ± 0.06 | 1.12 ± 0.24 | 0.3 | 5.4 |
| PpGPo1AlkB | F164I | 0.29 ± 0.03 | 1.54 ± 0.05 | 0.2 | 3.0 |
| PpGPo1AlkB | F164V | 0.28 ± 0.03 | 1.40 ± 0.19 | 0.2 | 3.2 |
| PpGPo1AlkB | L265W | 0.00 ± 0.00 | 0.11 ± 0.00 | 0.0 | 0.0 |
| PpGPo1AlkB | L265Y | 0.10 ± 0.01 | 0.75 ± 0.16 | 0.1 | 2.1 |
| PpGPo1AlkB | A268I | 0.00 ± 0.00 | 0.07 ± 0.01 | 0.0 | 0.0 |
| PpGPo1AlkB | A268F | 0.08 ± 0.01 | 0.08 ± 0.04 | 1.0 | 15.1 |
| PpGPo1AlkB | I233V+F164L | 0.87 ± 0.17 | 3.95 ± 0.39 | 0.2 | 3.5 |
| M_AlkB | WT | 0.64 ± 0.06 | 7.18 ± 0.35 | 0.1 | 1.0 |
| M_AlkB | I238V | 1.70 ± 0.05 | 11.92 ±1.14 | 0.1 | 1.6 |
| M_AlkB | F169L | 1.23 ± 0.07 | 4.38 ± 1.29 | 0.3 | 3.2 |
| M_AlkB | I238V+F169L | 0.72 ± 0.14 | 3.18 ± 0.36 | 0.2 | 2.5 |
| PpP1AlkB | WT | 0.48 ± 0.05 | 1.54 ± 0.01 | 0.3 | - |
| AboAlkB | WT | 0.41 ± 0.04 | 2.22 ± 0.87 | 0.2 | - |

The formation of an unknown side-product **3x** was observed for F164 variants (GPo1AlkB; F169 in M_AlkB) in GC-FID analysis (**Figure S7 A** and **B**). The product elutes directly after **3a**. The mass spectrogram of **3x** (**Figure S7 C**) shows structural similarity to the terminal alcohol **3a** (**Figure S5 B**) as well as to the substrate **2**, however more detailed analysis is needed to identify the product and understand the mechanisms behind its formation.

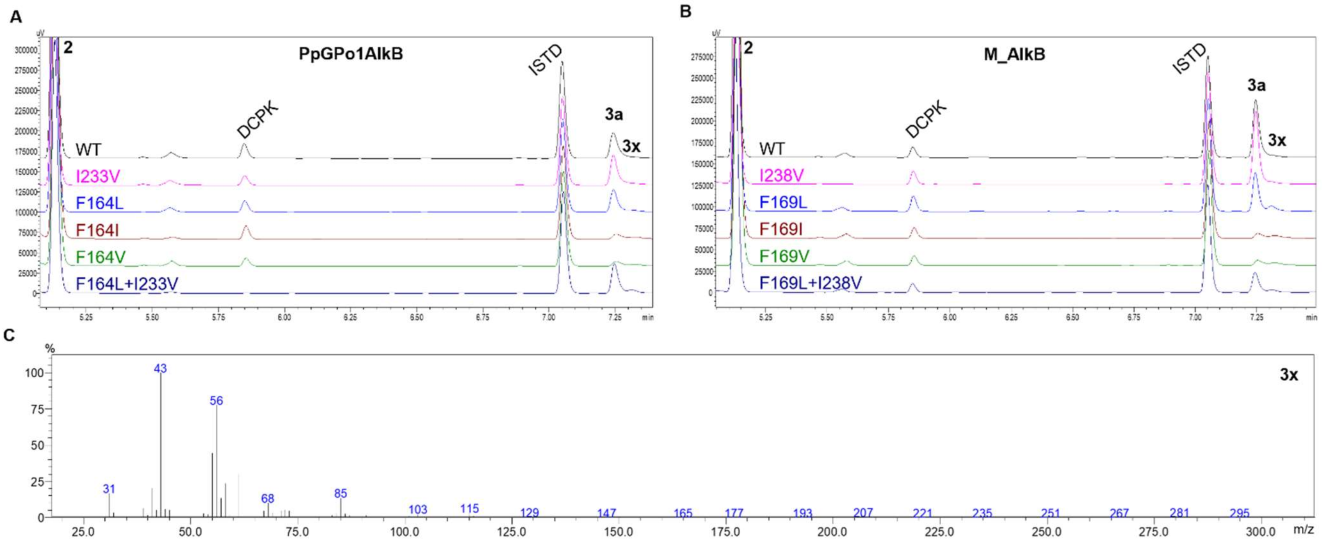

**Figure S7:** GC-FID analysis of 3 h reaction with variants of PpGPo1AlkB (**A**) and M_AlkB (**B**) and **2** as substrate revealed formation of an unidentified side-product **3x** for F164/F169 substitutions. The mass spectrum (**C**) of the side-product shows structural similarity to the substrate **2** and the main product **3a**.

#### 2.4 Whole-cell conversion of isoprenol by AlkBFGT

Conversion of isoprenol **1** to 2-methylene-1,4-butandiol (MBD) by AlkBFGT was also investigated to see if there would be any potential cross-reactions in whole-cell applications with the acetylation reaction of **1** to isoprenyl acetate **2**. The reaction was performed analogously to the whole-cell conversion of **2** using 5 mM of **1** as substrate. Samples for GC-MS analysis were taken directly at the beginning of the reaction, after 24 h and 48 h of reaction time and was extracted with EtoAc and dried over $NaSO_4$. The extract was derivatized using BSTFA as silylating agent. Peaks were assigned by comparison to commercially available standards. The silylated substrate **1** elutes at 6.05 min. The mono-silylated MBD derivates elute at around 8 min and the di-silylated MBD at 8.54 min. The reaction samples do not show any peaks which could be assigned to 2-methylene-1,4-butandiol, hence we assume that **1** is not accepted as substrate by AlkB and a cross-reaction with the acetylation reaction in a whole-cell procedure is not expected (**Figure S8**).

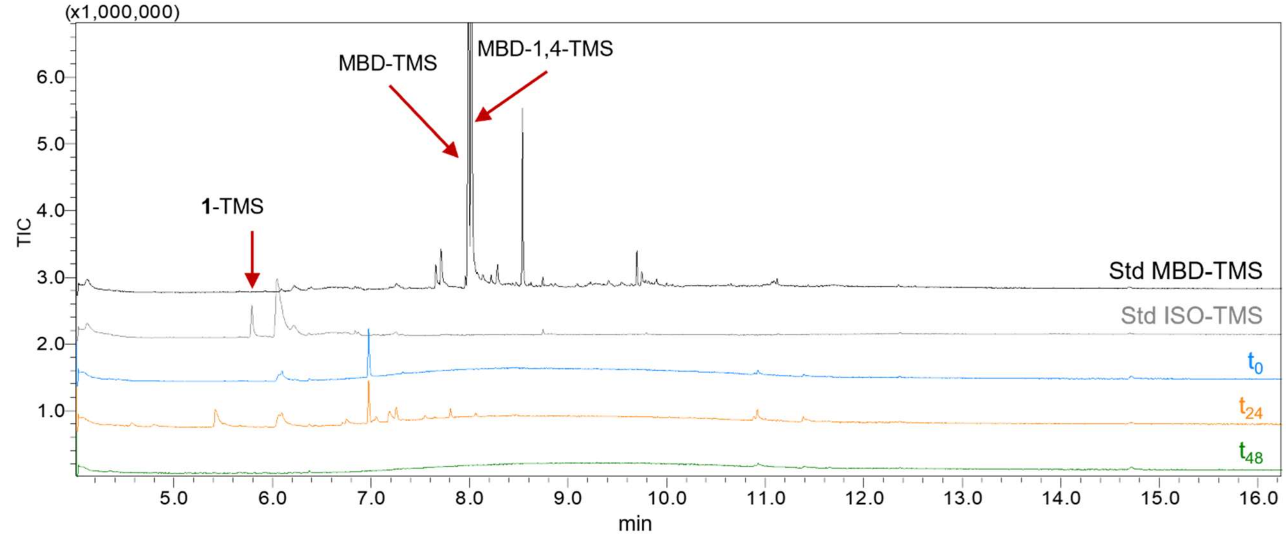

**Figure S8:** Comparison of GC-MS chromatograms of conversion of isoprenol **1** by AlkBFGT. The reaction was followed over time and samples were taken directly at the start of the reaction (blue), after 24 h (orange) and 48 h (green). The samples were derivatized using BSTFA. By comparing the reaction samples with the derivatized standards of **1** (grey) and MBD (black) formation of MBD by AlkBFGT from **1** can be excluded. Retention times at given conditions: 1-TMS: 6.05 min; MBD-1-TMS: 7.99 min; MBD-4-TMS: 8.02 min; MBD-1,4-TMS: 8.54 min. BSTFA: 5.79 min.

#### 2.5 Lactonization of 4-acetoxy-2-methylenebutyric acid to tulipalin A

The hydrolysis and lactonization of 4-acetoxy-2-methylenebutyric acid **5** to tulipalin A **6** was tested as a one-step reaction by incubation of **5** with concentrated HCl at 80 °C. Already after 3 h **5** was fully lactonized to **6** (**Figure S9**).

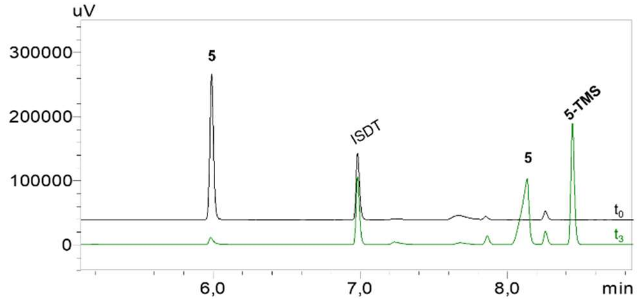

**Figure S9:** 4-acetoxy-2-methylene- butyric acid **5** can be fully converted to tulipalin A **6** within 3 h by acidification with concentrated HCl and incubation at 80°C (4c-TMS - BSTFA-derivatized 4c). Samples were extracted with EtoAc (1:10 (v/v)) containing 1 mM methyl benzoate as internal standard (ISTD) and analyzed with GC-FID.
